# Phenethylamine-producing gut bacteria induces diarrhea-predominant irritable bowel syndrome by increasing serotonin biosynthesis

**DOI:** 10.1101/2022.03.05.483096

**Authors:** Lixiang Zhai, Chunhua Huang, Ziwan Ning, Yijing Zhang, Min Zhuang, Wei Yang, Xiaolei Wang, Jingjing Wang, Eric Lu Zhang, Haitao Xiao, Ling Zhao, Yan Y. Lam, Chi Fung Willis Chow, Jiandong Huang, Shuofeng Yuan, Kui Ming Chan, Hoi Leong Xavier Wong, Zhao-xiang Bian

## Abstract

Despite the strong association between gut microbial dysbiosis, serotonin (5-HT) dysregulation and diarrhea-predominant irritable bowel syndrome (IBS-D), the mechanism by which changes in the gut microbiota contribute to the pathogenesis of IBS-D, particularly the role of dysregulated 5-HT production, remains unclear. The present study identified *Ruminococcus gnavus* in the human gut microbiota as a key risk factor of IBS-D. *R. gnavus* was significantly enriched in IBS-D patients and exhibited positive correlation with serum 5- HT level and severity of diarrhea symptoms. We showed that *R. gnavus* induced diarrhea-like symptoms in mice by promoting microbial shunting of essential aromatic amino acids to aromatic trace amines including phenethylamine and tryptamine, thereby stimulating the biosynthesis of peripheral 5-HT, a potent stimulator for gastrointestinal transit. This study identify gut-microbial metabolism of dietary amino acids as a cause of IBS-D and lays a foundation for developing novel therapeutic target for the treatment of IBS-D.

**Graphical abstract:** 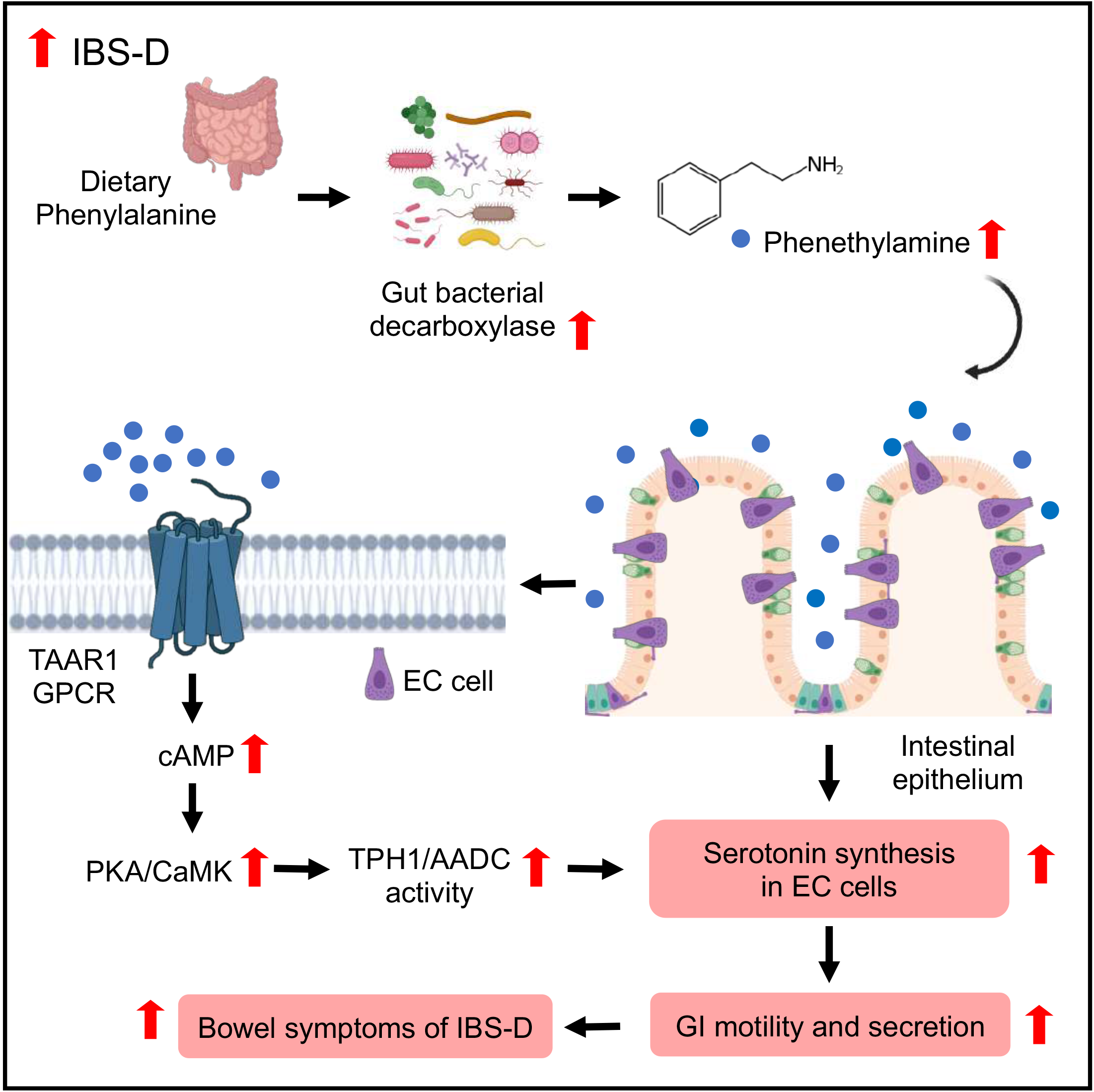

## Introduction

Irritable bowel syndrome (IBS) is one of the most prevalent functional bowel disorders characterized by an array of gastrointestinal (GI) symptoms including abdominal pain, bloating, abdominal distention and bowel habit abnormalities (Sperber et al., 2017). IBS is associated with comorbid conditions that substantially reduce the quality of life, leading to a growing social and economic burden worldwide. Despite the high prevalence of IBS, treatment options for the management of IBS are limited (Van den Houte et al., 2020), and most therapeutic approaches can only relieve individual symptoms of IBS instead of curing it..

Serotonin (5-HT) is an important neurotransmitter synthesized from tryptophan by tryptophan decarboxylase 1 (TPH1) in intestinal enterochromaffin cells (ECs) and tryptophan decarboxylase 2 (TPH2) in enteric and central serotonergic neurons (Bellono et al., 2017). 5-HT released from EC cells in the GI tract modulates gut motility and hypersensitivity functions (Kendig and Grider, 2015). EC cell hyperplasia and increased 5-HT production are often observed in the colon of IBS patients, especially those with diarrhea-predominant IBS (IBS-D) (Dunlop et al., 2003; Thijssen et al., 2016). Increased production of peripheral 5-HT from intestinal EC cells can lead to intestinal symptoms in IBS-D (Wong et al., 2019). Therapeutic interventions targeting various 5-HT receptors have been shown to be effective in the management of IBS, but their therapeutic effects are hampered by adverse complications, such as ischemic colitis and severe constipation (Stasi et al., 2014). Therefore, an improved understanding of the mechanism underlying increased 5-HT production in IBS-D patients may pave the way for innovative therapeutic strategies against IBS-D.

Emerging evidence reveals that gut microbiota plays an important role in the pathophysiology of IBS. In alignment with other studies, our previous findings have reported significant changes in the structure of gut microbiota in patients with IBS, especially in IBS-D (Han et al., 2021; Jeffery et al., 2020; Zhao et al., 2020). Transplantation of fecal microbiota from IBS-D patients results in IBS-like symptoms including accelerated GI transit and intestinal barrier dysfunction in recipient germ-free mice (De Palma et al., 2017). Despite the strong association between gut dysbiosis and the pathogenesis of IBS, the mechanism by which changes in the gut microbiota contributes to the development of IBS remain unclear.

Gut microbiota is essential for the maintenance of homeostatic peripheral 5-HT levels in the host since serum 5-HT levels are remarkably reduced in germ-free mice (Reigstad et al., 2015; Yano et al., 2015). Moreover, gut microbial metabolites, such as secondary bile acids and short-chain fatty acids, stimulate peripheral 5-HT biosynthesis and enhances GI motility in mice (Ge et al., 2018). However, the identity of bacterial species contributing to the regulation of peripheral serotonin production, the mechanism of action of their serotonin-stimulating metabolites and their pathophysiological roles in IBS-D remain to be elucidated. To inform intervention strategies for the treatment of IBS-D, it is imperative to characterize gut bacteria and identify gut-microbial metabolites that regulate peripheral 5-HT levels in IBS-D patients.

## Results

### The positive association between *Ruminococcus gnavus*, peripheral serotonin and severity of diarrhea symptoms in IBS-D

To understand the role of gut microbiota responsible for the deregulation of peripheral 5-HT in the pathophysiology of IBS-D, a total of 290 IBS-D patients and 89 healthy controls (HC) were recruited in our previous study (Zhao et al., 2019) and their biospecimens including feces, serum and urine were used for analyses. We firstly analysed the serum 5-HT and urine 5-HIAA (a urinary biomarker of 5-HT) levels in this cohort. Consistent with our previous and other studies (Thijssen et al., 2016; Wong et al., 2019), increased level of peripheral serotonin (5-HT) was found in the sera from IBS-D patients (Figure.1A). In line with elevated serum 5-HT level, increased level of urine 5-HIAA was observed in IBS-D patients (Figure.1B). To identify the gut bacteria that were potentially responsible for the dysregulation of peripheral 5-HT, we performed correlation analysis between gut microbiota and serum 5-HT level in IBS-D patients. Notably, we found a series of bacteria species positively correlated with serum 5-HT level in IBS-D patients (Figure.1C and Supplement Table.S1). Among these bacteria species, *Ruminococcus gnavus*, a culturable gut bacterium with relatively higher abundance in human gut microbiota, was significantly increased in IBS-D patients (Figure.1D) and positively correlated with serum 5-HT level in IBS-D patients (Figure.1E). Furthermore, *R. gnavus* abundances were positively correlated with severity of IBS-D symptoms assessed by Bristol stool scale in IBS-D patients (Figure.1F). These data collectively showed that significant changes of bacteria species, particularly *R. gnavus*, may be associated with altered 5-HT metabolism in IBS-D.

**Figure 1.**
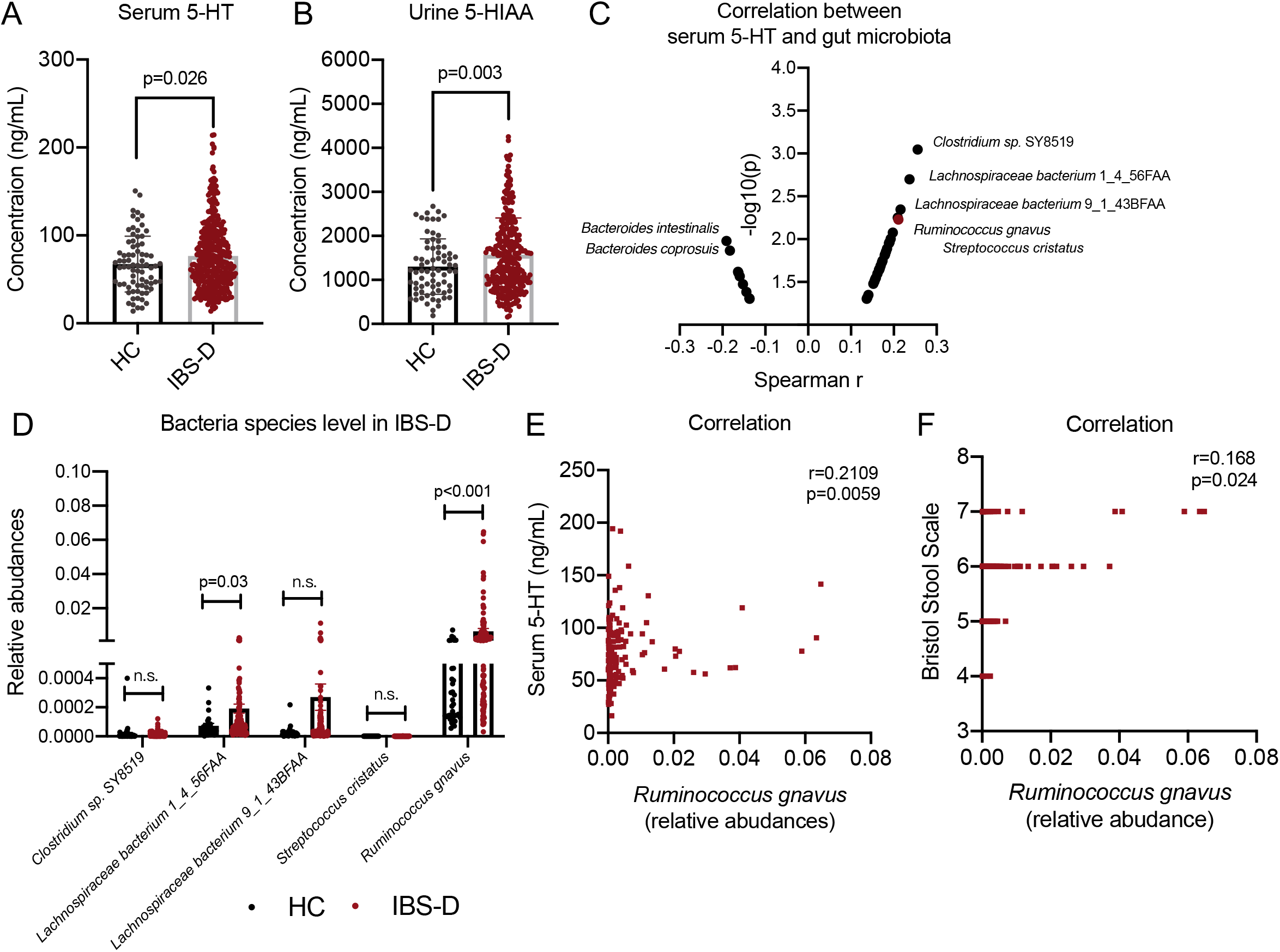
*R. gnavus* significantly increased in IBS-D patients and positively correlated with serum 5-HT level and diarrhea symptoms. (A-B) Serum 5-HT and urine 5-HIAA level in HC (n=89) and IBS-D (n=290) subjects. (C) Spearman r value and p value of gut bacteria species abundances and serum 5-HT level in IBS-D patients. (D) Relative abundances of selected gut bacteria species in HC and IBS-D subjects. (E) Spearman’s correlation between relative abundances of *R. gnavus* with serum 5-HT level in IBS-D subjects. (F) Spearman’s correlation between relative abundances of *R. gnavus* with Bristol Stool Scale in IBS-D subjects. Differences of relative abundances of *R. gnavus* in HC and IBS-D patients were analyzed by one-tailed Mann-Whitney test. Data are presented as mean + S.E.M.

### Monoassociation with *Ruminococcus gnavus* stimulated serotonin production, induced diarrhea-like symptoms in accompanied with phenethylamine production

To investigate the pathological role of *R. gnavus* in dysregulated 5-HT production in IBS-D, we monoassociated pseudo-germ-free mice with a commercially available gut bacteria strain *Ruminococcus gnavus* (ATCC 29149). Pseudo germ-free mice monoassociated with *R. gnavus* exhibited significantly elevated levels of serum and intestine 5-HT (Figure.2A-B). In line with the increased 5-HT level in serum and intestine, shortened GI transit time and increased fecal water content were observed in pseudo germ-free mice monoassociated with *R. gnavus* (Figure.2C-D). These results demonstrated that monoassociation with *R. gnavus* led to increased 5-HT production and IBS-D-like symptoms in mice.

**Figure 2.**
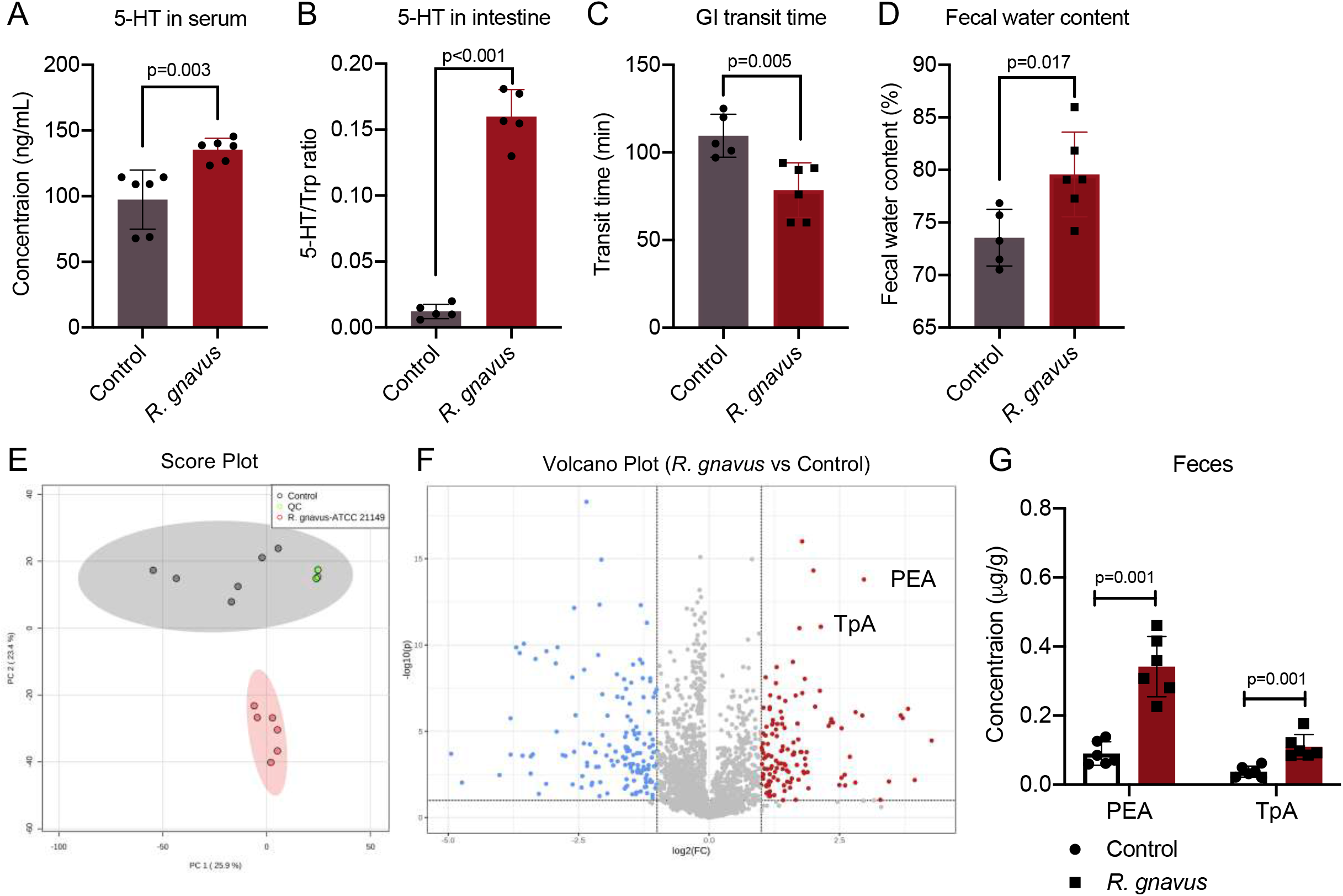
*R. gnavus* induces elevated 5-HT production and diarrhea-like symptoms in accompanied with PEA and TpA production in mice. (A) Serum and intestine 5-HT level in pseudo germ-free mice after monoassocation with/without *R. gnavus* (ATCC 29149) (n=6/group). (C-D) GI transit time and fecal water content indexes in pseudo germ-free mice after monoassocation with/without *R. gnavus* (ATCC 29149) (n=6/group). (E) Score plot of fecal metabolome in pseudo germ-free mice after monoassocation with/without *R. gnavus* (ATCC 29149) (n=6/group). (F) Volcano plot of fecal metabolome in pseudo germ-free mice after monoassocation with/without *R. gnavus* (ATCC 29149) (n=6/group). (G) Fecal PEA and TpA level in pseudo germ-free mice after monoassocation with/without *R. gnavus* (ATCC 29149) (n=6/group). Differences of serum and intestine 5-HT level and GI transit time and fecal water content indexes in mice were analyzed by two-tailed student t-test. Differences of fecal PEA and TpA level in mice were analyzed by one-tailed student t-test. Data are presented as mean + S.D. See also Supplement Figure.S1.

To further investigate the molecular mechanisms underlying the microbial regulation of 5-HT production, we performed untargeted metabolomics to identify the changes of fecal metabolome in pseudo germ-free mice with/without monoassociation with *R. gnavus*. Score and volcano plots showed dramatic differences in metabolic profiles of fecal samples between the two groups (Figure.2E and Figure.2F). Notably, a significant elevation of aromatic trace amines including PEA and TpA was detected in fecal samples of mice monocolonized with *R. gnavus* (Figure.2F-G). Moreover, the abilities of *R. gnavus* in converting Phe and Trp into PEA and TpA respectively were validated *in vitro* (Supplement Figure.S1A-B). These data showed PEA and TpA may be associated with the stimulatory effects of *R. gnavus* on 5-HT production and GI transit.

### The positive association between phenethylamine and peripheral serotonin in IBS-D

To investigate the clinical relevance of our findings obtained from mouse studies, we examined PEA and TpA levels in our IBS-D patient cohort. Consistently, we found that PEA was significantly increased in the faeces of IBS-D patients (Figure.3A). In contrast, there was no significant change in phenylalanine (Phe), the precursor of PEA, in the faeces of IBS-D patients (Figure.3B). Notably, correlation analysis revealed that fecal PEA level was positively correlated with *R. gnavus* at a highest r value and *p* value among other gut bacterial species in IBS-D patients (Figure.3C and Supplement Table.S2). Furthermore, PEA level was also positively correlated with the severity of diarrheal symptoms measured by Bristol stool scale and serum 5-HT level in IBS-D patients (Figure.3D-F). In line with the elevated level of PEA in IBS-D patients, faecal TpA level, but not Trp level, was also found significantly increased in IBS-D patients (Supplement Figure.S2A-B) and positively correlated with Bristol stool scale and serum 5-HT level in IBS-D patients (Supplement Figure.S2C-D). We then compared the catalytic ability of gut microbiota from HC and IBS-D in transforming Phe into PEA as well as Trp into TpA by batch culture using faecal samples *in vitro*. Higher concentrations of PEA and TpA were detected in the culture medium Supplemented with the bacterial suspension of IBS-D fecal samples (Figure.3G and Supplement Figure.S2E), indicative of high catalytic ability of gut microbiota in processing Phe and Trp into PEA and TpA in IBS-D patients. These data collectively showed that the changes of fecal PEA and TpA are positively associated with peripheral 5-HT and severity of diarrheal symptoms in IBS-D.

**Figure 3.**
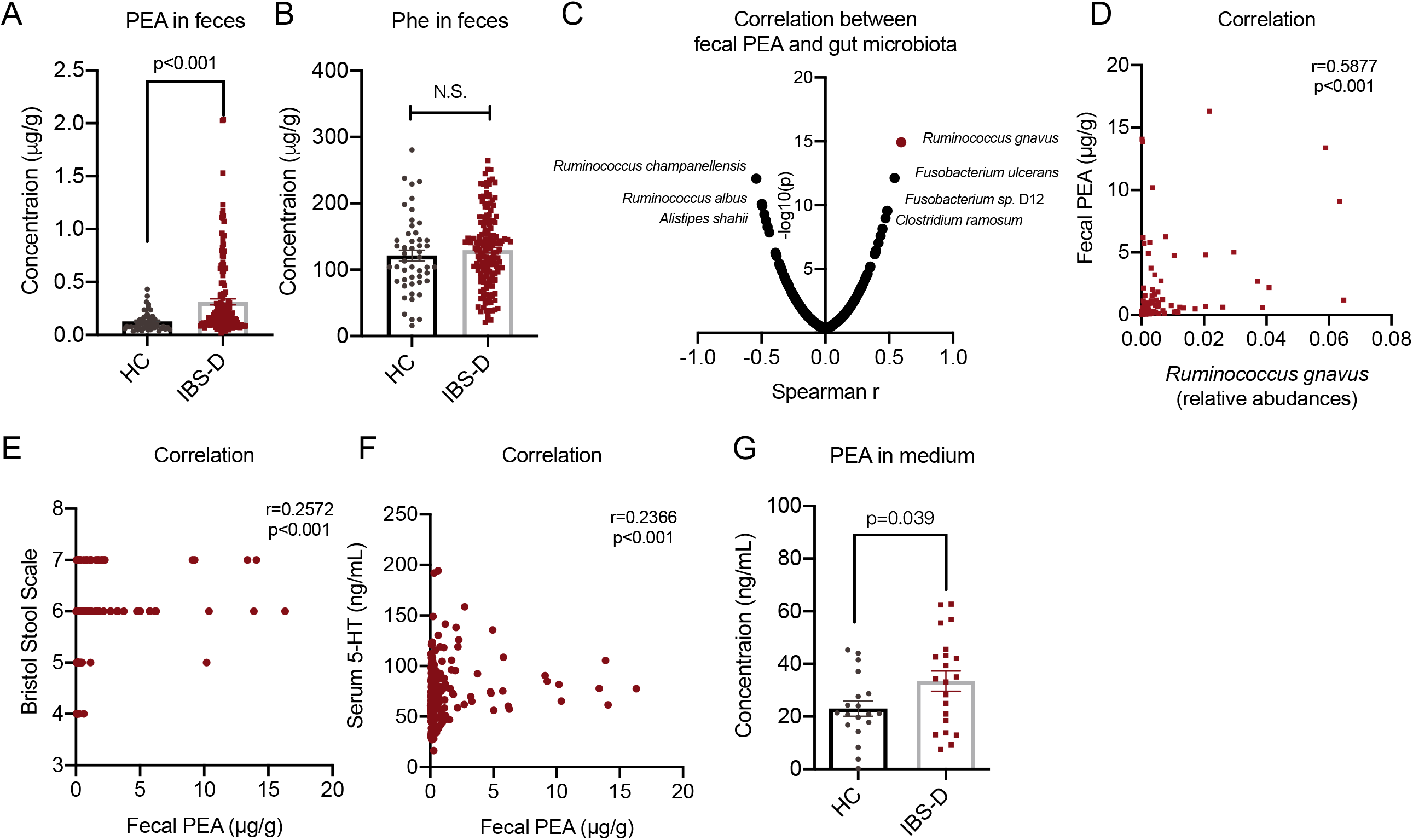
Fecal PEA and TpA are increased and positively correlated with serum 5-HT level in IBS-D patients. (A-B) PEA and Phe level in fecal samples of in HC (n=89) and IBS-D (n=290) subjects. (C) Spearman r value and p value of gut bacteria species abundances and fecal PEA level in IBS-D patients. (D-F) Spearman’s correlation between fecal PEA level with *R. gnavus* abudances, Bristol Stool Scale and serum 5-HT level in IBS-D subjects. (G) PEA level in batch culture samples using feces from HC and IBS-D (n=30/group). Differences of fecal PEA and Phe level in HC and IBS subjects as well as batch culture samples were analyzed by one-tailed Mann-Whitney test. Data are presented as mean + S.E.M. See also Supplement Figure.S2.

### Phenethylamine accelerates gastrointestinal motility by stimulating serotonin biosynthesis

Since fecal PEA level and TpA level was significantly increased in IBS-D patients and positively correlated with peripheral 5-HT along with diarrhea-related symptoms in IBS-D patients, we postulated that PEA and TpA stimulate 5-HT production and hence regulate the GI transit and colonic secretion. To address this hypothesis, we examined the effects of PEA and TpA on 5-HT production in mouse intestinal tissues *ex vivo* and intestinal organoids *in vitro*. Notably, PEA treatment within the range of pathophysiological concentrations detected in IBS-D patients significantly enhanced the 5-HT production in a dose-dependent manner in both mouse intestinal tissues and intestinal organoids (Figure.4A-B). Similar observations on the stimulatory effect of PEA on 5-HT production were found in QGP-1 cells, a well-established human pancreatic endocrine cell line used for studying 5-HT production (Supplement Figure.S3A-C). In contrast, treatment with phenylalanine (Phe), the PEA precursor, and phenylacetic acid (PA), the downstream metabolite of PEA, did not change the 5-HT level in QGP-1 cells (Supplement Figure.S3D). We then investigated whether PEA regulates GI transit and colonic secretion in mice by modulating 5-HT production. Notably, treatment with PEA within the pathophysiological concentrations by oral administration resulted in significantly enhanced GI transit and fecal water content along with elevated levels of serum and intestinal 5-HT in mice (Figure.4C-F). Blockade of 5-HT production by TPH1 inhibitor LX-1031 did not only effectively inhibited PEA-induced increase in 5-HT levels (Figure.4G-H), but also completely suppressed the increased GI transit and fecal water content in PEA-treated mice (Figure.4I-J). Similarly, we found that TpA exerted stimulatory effects on intestinal 5-HT production *ex vivo*, *in vivo* and *in vitro* (Supplement Figure.S3E-G). In contrast, 5-HT production was not altered by the treatment with precursor (Trp) or metabolite (IAA) of TpA (Supplement Figure.S3H). Interestingly, we noticed TpA and PEA exhibit additive effects on 5-HT production in intestine tissues cultured *ex vivo* (Supplement Figure.S3 I). Collectively, these results demonstrated that PEA and TpA stimulate GI transit and secretion in a 5-HT-dependent manner.

**Figure 4.**
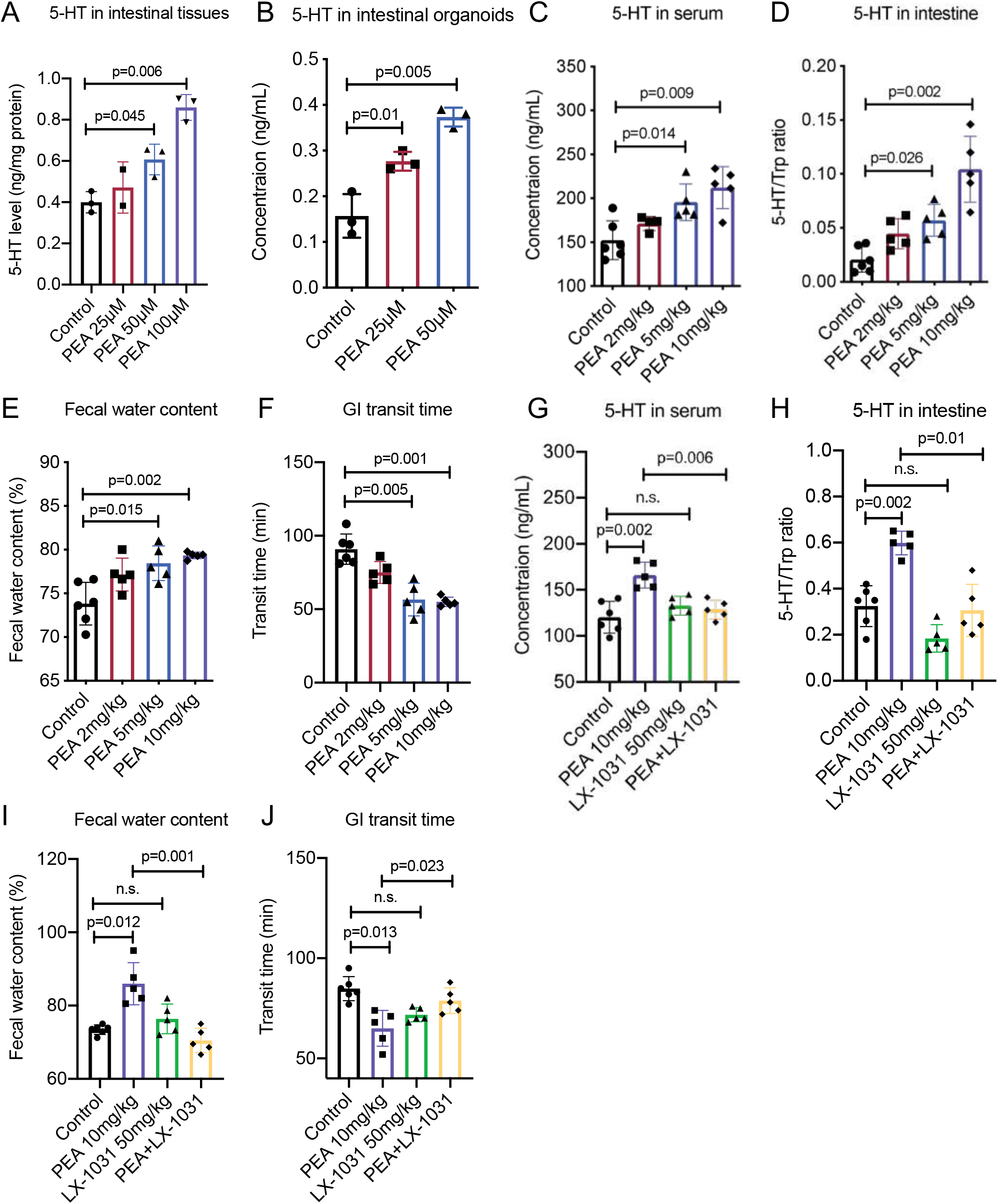
PEA and TpA activates 5-HT production, accelerates GI transit and increase colonic secretion *in vitro* and *in vivo*. (A) 5-HT level in mice *ex vivo* intestinal tissues after treatment of PEA as indicated concentration (25μM, 50μM and 100μM) or control for 2 hours (n=3/group). (B) 5-HT level in mice *in vitro* intestinal organoids after treatment of PEA as indicated concentration (25μM and 50μM) or control for 4 hours (n=3/group). (C-D) 5-HT level in mice serum and intestinal tissues after treatment of PEA as indicated dosages (2mg/kg, 5mg/kg and 10mg/kg) or control (water) (n=6/group). (E-F) Fecal water content and GI transit time in mice after treatment of PEA as indicated dosages (2mg/kg, 5mg/kg and 10mg/kg) or control (water) (n=6/group). (G-H) 5-HT level in mice serum and intestinal tissues after treatment of PEA (10mg/kg), TPH1 inhibitor (LX-1031, 50mg/kg) or control (water) (n=6/group). (I-J) Fecal water content and GI transit time in mice after treatment of PEA (10mg/kg) and TPH1 inhibitor (LX-1031, 50mg/kg) or control (water) (n=6/group). Differences of 5-HT level, GI transit time and fecal water content were analyzed by t-test (two-tailed Mann-Whitney test) or ordinary one-way ANOVA. Data are presented as mean + S.D. See also Supplement Figure.S3.

### Phenethylamine stimulates serotonin production via a TAAR1 dependent mechanism

The peripheral 5-HT is synthesized from Trp by TPH1 (Trp to 5-HTP) and AADC (5-HTP to 5-HT) and subsequently metabolized into 5-HIAA by MAO/ALDH (Matthes and Bader, 2018). To further investigate the mechanism underlying the stimulatory effects of PEA on 5-HT production, we analysed the relative ratios of 5-HTP/Trp, 5-HT/5-HTP and 5-HIAA/5-HT by determining Trp, 5-HTP and 5-HIAA levels in serum of PEA-treated mice (Supplement Figure.S4A-C). In addition to the upregulation of 5-HT, we found that PEA treatment in a dose-dependent manner also led to the increase in 5-HTP level in the serum of mice. As a result, 5-HTP/Trp and 5-HT/5-HTP ratios, but not 5-HIAA/5-HT ratio, were increased in serum of PEA-treated mice (Figure.5A), suggesting the stimulatory effect of PEA on 5-HT signaling is likely mediated by biosynthetic pathway (TPH1 and AADC) but independent of catabolic pathway (MAO/ALDH).

**Figure 5.**
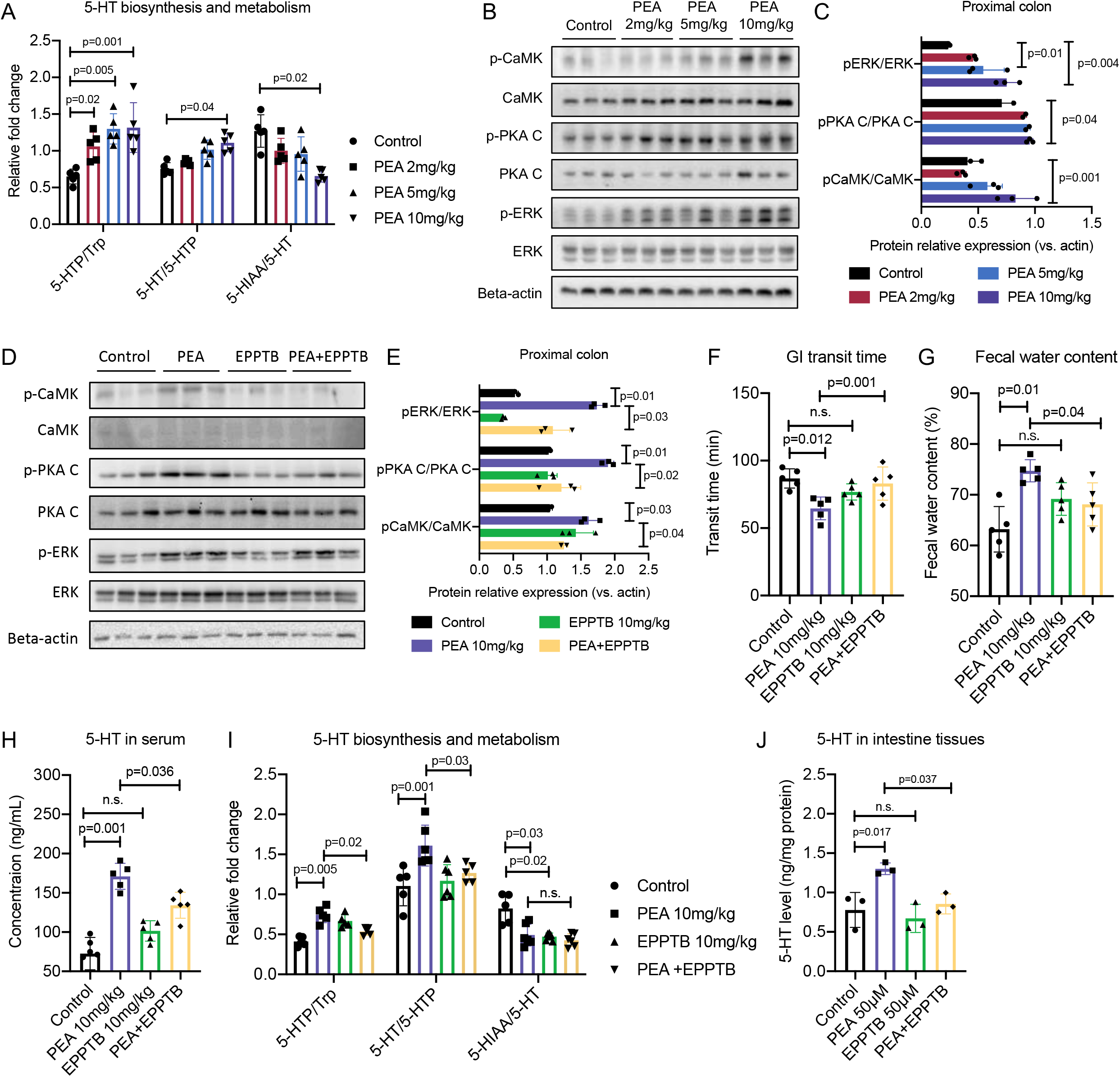
PEA stimulates 5-HT production via a TAAR1 dependent mechanism. (A) 5-HT biosynthesis and metabolism profiles after treatment of PEA as indicated dosages (2mg/kg, 5mg/kg and 10mg/kg) or control (water) (n=6/group). (B-C) Western blot (and semi-quantification) in proximal colonic tissues of mice after treatment of PEA as indicated dosages (2mg/kg, 5mg/kg and 10mg/kg) or control (water) (n=3/group). (D-E) Western blot (and semi-quantification) in proximal colonic tissues of mice after treatment of PEA (10mg/kg) and TAAR1 antagonist EPPTB (10mg/kg) or control (1% DMSO in saline) (n=3/group). (F-G) GI transit time and fecal water content in mice after treatment of PEA (10mg/kg) and TAAR1 antagonist EPPTB (10mg/kg) or control (1% DMSO in saline) (n=6/group). (H) 5-HT level in mice serum after treatment of of PEA (10mg/kg) and TAAR1 antagonist EPPTB (10mg/kg) or control (1% DMSO in saline) (n=6/group). (I) 5-HT biosynthesis and metabolism profiles after treatment of PEA (10mg/kg) and TAAR1 antagonist EPPTB (10mg/kg) or control (1% DMSO in saline) (n=6/group). (J) 5-HT level in mice *in vitro* intestinal tissues after treatment of PEA (50μM) and TAAR1 antagonist EPPTB (50μM) or control (1% DMSO) (n=3/group). Differences of 5-HT level, GI transit time and fecal water content were analyzed using ordinary one-way ANOVA. Data are presented as mean + S.D. See also Supplement Figure.S4.

Trace amine-associated receptor 1 (TAAR1), a G protein-coupled receptor (GPCR), is a well-known receptor for PEA (Xie and Miller, 2008). Previous studies revealed that enzymes responsible for biosynthesis of 5-HT, including TPH1 and AADC, are activated by downstream mediators of GPCR signaling, namely cyclic AMP-dependent protein kinase A (PKA) and calcium/calmodulin-dependent kinase (CaMKII) (Duchemin et al., 2000; Kuhn et al., 1997; Kumer et al., 1997; Neff et al., 2002; Young et al., 1993). We found that PEA indeed activated PKA and CaMKII, as indicated by the increased phosphorylation of these proteins in colonic tissues of mice treated with PEA (Figure.5B-C). PEA-induced activation of PKA and CaMKII was abolished by inhibition of TAAR1 with specific antagonist EPPTB (Figure.5D-E). Consistently, blockade of TAAR1 activities by EPPTB also suppressed PEA-induced 5-HT production, GI transit and colonic secretion in mice (Figure.5F-J) as well as 5-HT elevation in intestine tissues cultured *ex vivo*. In line with these findings, the PEA-induced reduction in the ratios of 5-HTP/Trp and 5-HT/5-HTP in serum was also abrogated by EPPTB treatment (Figure.5I and Supplement Figure.S4D-G). These results suggest that PEA acting through TAAR1 promotes GI transit and increases colonic secretion by stimulating 5-HT production.

### Phenethylamine produced by IBS-D associated gut microbiota enhances serotonin synthesis and gastrointestinal transit *in vivo*

To study *in vivo* action of PEA on 5-HT production and GI motility, we integrated the plasmid expressing tryptophan decarboxylase sequence (TDC), an enzyme that catalyses the conversion of Phe into PEA and Trp into TpA from *R. gnavus* (strain ATCC 29149), into a gut microbe *E.coli* K12. Pseudo germ-free mice were colonized with either vector-control *E. coli* or *E. coli* TDC^*+*^ by oral gavage. The successful integration of the plasmid was validated by PCR analyses and on *in vitro* production of PEA and TpA in LB medium assessed by LC-MS analyses (Supplement Figure.S5A-C). Pseudo germ-free mice colonized with *E. coli* TDC^*+*^ exhibited significantly elevated levels of PEA and TpA in faeces, confirming that *E. coli* TDC^*+*^ strain produced PEA *in vivo* (Figure.6A). Consistently, increased serum 5-HT level and fecal water content coupled with shortened GI transit time were observed in mice colonized with *E. coli* TDC^*+*^ compared with mice with vector-control *E. coli* (Figure.6B-D and Supplement Figure.S5D-E). These results demonstrated engineered bacteria with TDC produce PEA and TpA *in vivo* to stimulate 5-HT production, leading to accelerated GI transit and increased colonic secretion. To further investigate whether PEA and TpA-mediated TAAR1 signaling is involved in the elevation of 5-HT production and diarrhea-like symptoms induced by *R. gnavus*, we used TAAR1 antagonist EPPTB to block the action of PEA and TpA on 5-HT production. We showed that *R. gnavus-*induced 5-HT elevation in serum and intestine, as well as diarrhea-like symptoms including accelerated GI transit and increased fecal water content, were significantly suppressed by EPPTB treatment (Figure.6E-H and Supplement Figure.S5F-G). These results demonstrated aromatic trace amines-producing gut bacteria *R. gnavus* modulates 5-HT production and hence GI transits via PEA/TpA-mediated TAAR1 signalling, suggesting PEA-producing gut bacteria plays an important role in the pathogenesis of diarrhea symptoms of IBS-D patients.

**Figure 6.**
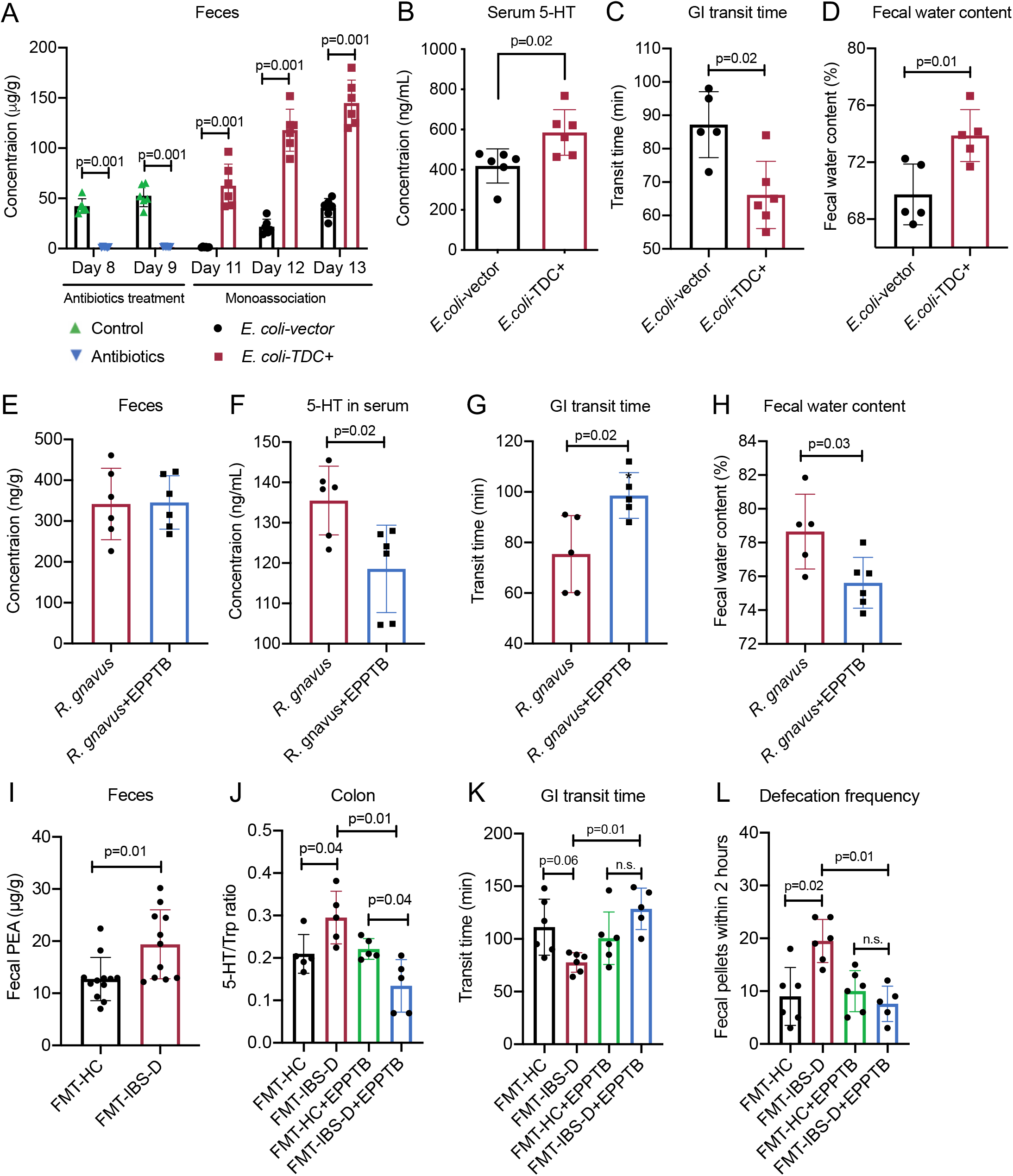
*In vivo* PEA and TpA produced by IBS-D associated bacteria enhances 5-HT synthesis and induce diarrhea-like symptoms. (A) Fecal PEA level in pseudo germ-free mice after monoassocation with *E.coli* vector control or *E.coli* TDC^+^ (n=6/group). (B-D) Serum 5- HT level, GI transit time and fecal water content in pseudo germ-free mice after monoassocation with *E.coli* vector control or *E.coli* TDC^+^ (n=6/group). (E-H) Fecal PEA level, serum 5-HT level, GI transit time and fecal water content in pseudo germ-free mice after monoassocation with *R. gnavus* (ATCC 29149) and TAAR1 antagonist EPPTB (10mg/kg) or control (1% DMSO in saline) (n=6/group). (I-L) Fecal PEA level, colon 5-HT level, GI transit time and defecation frequency in pseudo germ-free mice after monoassocation with gut microbiota from HC (n=8) and IBS-D (n=8) subjects and TAAR1 antagonist EPPTB (10mg/kg) or control (1% DMSO in saline) (n=6/group). Differences of PEA level, 5-HT level in serum and intestine, GI transit time defecation frequency were analyzed using student t-test (two-tailed Mann-Whitney test) or ordinary one-way ANOVA. Data are presented as mean + S.D. See also Supplement Figure.S5.

To investigate the therapeutic potential of targeting TAAR1 in the management of IBS-D, we made use of the mice colonized with gut microbiota derived from IBS-D patients or healthy control as a preclinical model of IBS-D and treat them with TAAR1 antagonist EPPTB. Pseudo germ-free mice colonized with IBS-D microbiota exhibited diarrhea-like symptoms characterized by increased GI transit and defecation frequency, coupled with elevated 5HT biosynthesis and increased production of PEA and TpA in gut. All of these pathological changes induced by transplantation of IBS-D faecal microbiota were significantly suppressed by inhibition of TAAR1 activity with EPPTB. These data highlights the therapeutic potential of targeting PEA/TAAR1 pathway in the management of IBS-D.

## Discussion

In the present study, we provided mechanistic insights into the contribution of gut microbiota and its metabolites derived from dietary nutrients to the development of IBS-D by regulating 5-HT production. We herein showed human gut bacterium *R. gnavus* enriched in IBS-D patients is positively associated with peripheral 5-HT and severity of diarrheal symptoms. Monoassociation with *R. gnavus* in pseudo germ-free mice stimulated peripheral 5-HT production and hence induced IBS-D-like symptoms including accelerated GI transit and increased colonic secretion. Furthermore, we showed PEA and TpA produced by *R. gnavus-* mediated catabolism of dietary essential amino acids, are responsible for induction of diarrhea-like symptoms via the stimulation of 5-HT production. To the best of our knowledge, we are the first to show that PEA and TpA are the direct stimulators of 5-HT biosynthesis, thereby regulating GI transit and colonic secretion *in vivo* and *ex vivo*. Mechanistically,PEA and TpA bind to and activate their G-protein coupled receptor (TAAR1) which in turn mediates 5-HT biosynthesis by stimulating TPH1/AADC activities. To further validate our observations obtained from *in vitro* studies, we showed that manipulation of endogenous aromatic trace amines level by colonization of engineered bacteria with TDC led to IBS-D-like diarrheal symptoms coupled with enhanced 5-HT production. Collectively, these findings suggest PEA and TpA producers (eg. *R. gnavus*) stimulate 5-HT biosynthesis to accelerate GI transit via a TAAR1-dependent mechanism, thereafter contributing to diarrheal symptoms of IBS-D patients. This study demonstrates the causality between gut dysbiosis and 5-HT-associated diarrheal symptoms and confirms the dysregulation of host-microbe interaction as one of the leading causes of IBS-D.

Although gut microbiota has been shown to modulate peripheral 5-HT and GI motility by interacting with host EC cells (Agus et al., 2018), the mechanism by which gut-microbial metabolites affect 5-HT production and their roles in the development of IBS remain largely unclear. Gut microbial metabolites including butyrate, propionate, tyramine, deoxycholate and p-aminobenzoate have been shown to stimulate 5-HT biosynthesis *in vitro*, which provide fundamental understanding to the regulatory mechanisms of gut microbiota in regulating 5-HT (Yano et al., 2015). However, these metabolites including short fatty acids (butyrate and propionate) and deoxycholate are not altered in IBS-D patients (Luo et al., 2021; Wei et al., 2020). Although tyramine was found increased in inflammatory bowel disease (Santoru et al., 2017) and colorectal cancer (Salahshouri et al., 2021), we could not detect significant changes of tyramine and tyrosine in stool samples and fecal batch culture from IBS-D patients (Supplement Figure.S2 F-H). These data suggest that these metabolites may not contribute to the pathogenesis of IBS-D.

In this study, we further confirmed the causal relationship between gut microbiota dysbiosis and 5-HT dysregulation and their contribution to the pathogenesis of IBS-D. We for the first time identified human gut bacteria *R. gnavus* as a strong modulator of peripheral 5-HT level. Metabolically, PEA and TpA produced by *R. gnavus* exert potent stimulatory effects on 5-HT biosynthesis in intestinal EC cells. PEA, TpA and tyramine are aromatic trace amines generated from microbial metabolism of dietary aromatic amino acids in the host gut (Liu et al., 2020). In contrast, their precursors and downstream metabolites do not affect 5-HT production *in vitro* (Supplement Figure.S3D/H), suggesting that the regulation of 5-HT production by microbial breakdown of dietary aromatic amino acids is specifically mediated by aromatic trace amines. Previous studies showed that TpA drives fluid secretion and alters GI transit (Bhattarai et al., 2018), and tyramine exhibits direct stimulatory effects on intestine contraction (Marcobal et al., 2012). Therefore, aromatic trace amines control the GI motility by simultaneously regulating 5-HT production and its action, potentially explaining why the efficacy of current pharmacological approaches for treating IBS-D by targeting serotonin receptor is not satisfactory owing to the continuous activation of 5-HT biosynthesis by aromatic trace amines with the supply from dietary amino acids.

The elevation of TpA and its precursor Trp was detected in stool samples of IBS-D patients (Mars et al., 2020), and its action on GI motility via activating epithelial 5-HTR4 has been reported (Bhattarai et al., 2018). In contrast, the 5-HTR4 antagonist, which has been shown to block the TpA stimulatory action on GI transit (Bhattarai et al., 2018), failed to block the inducing effects of TpA and PEA on 5-HT production (Supplement Figure.S4 H). Therefore, developing new strategies to reduce microbial transformation of dietary amino acids into aromatic trace amines may pave the way towards precise treatment and eventually the cure of IBS-D. Gut microbial-produced aromatic trace mines including PEA, TpA and tyramine are ligands of TAAR1, a G-protein coupled receptor expressed in both central nervous system and gut (Sotnikova et al., 2009). Dysregulated TAAR1 signaling has been found to be involved in psychiatric disorders (Dodd et al., 2021) and mood disorders (Alnefeesi et al., 2021). TAAR1 modulators (agonists) are being studied as novel drugs for schizophrenia, Parkinson’s related psychoses and substance abuse (Dodd et al., 2021). Interestingly, about 60% of patients with neuropsychiatric disorders present GI symptoms, such as IBS (Fadgyas-Stanculete et al., 2014). Deregulation of TAAR1 ligands may therefore be a common factor in both GI disorders and comorbid neuropsychiatric disorders which can be addressed in future studies. The present study identified *R. gnavus* as an aromatic trace amines-producer that stimulates 5-HT production and induces IBS-D-like symptoms in mice. In other studies, *R. gnavus* has been reported to be associated with inflammatory bowel disease (IBD) (Hall et al., 2017) and exhibits proinflammatory properties by producing inflammatory polysaccharides (Henke et al., 2019). In line with these observations, low-grade chronic inflammation has been found in the colonic tissues of IBS-D patients (Öhman and Simrén, 2010; Rana et al., 2013). These observations suggest that *R. gnavus* may promote inflammatory responses and impair barrier functions to induce other IBS-D symptoms, such as abdominal pain and bloating, in addition to increased GI transit and secretion. In fact, studies has demonstrated that IBS patients indeed are at a greater risk of developing IBD (Porter et al., 2012). Our findings reveal that mice monoassociated with *R. gnavus* exhibited increased GI transit and colonic secretion without the presence of fecal occult blood, a major symptom of IBD, suggesting that *R. gnavus* predominantly contributes to the development of IBS-D rather than IBD in normal mice but may promote colonic injury in the experimental model of colitis.

Collectively, our findings not only provide new insights into the pathogenesis of IBS-D, but also lay a foundation for developing therapeutics for the management of 5-HT abnormalities in IBS-D. There are several limitations in this study. Due to lack of genetic tools to target the TDC gene in *R. gnavus*, we used TAAR1 antagonist to determine the effects of *R. gnavus* ATCC 29149 on GI motility and 5-HT production. Our investigation demonstrated that TAAR1 antagonist effectively abolishes the stimulatory effect of PEA on serotonin biosynthesis *in vitro* and *in vivo*, suggesting that the *in vivo* action of PEA is primarily mediated via TAAR1 in the context of gut motility. These studies highlight the therapeutic potential of targeting TAAR1 or microbial TDC enzyme in the treatment of IBS-D. However, other receptors, such as 5-HT4R, may also mediate the actions of aromatic trace amines such as TpA (Bhattarai et al., 2018), which should be addressed in the future studies.

## Abbreviations

5-HIAA: 5-Hydroxyindole acetic acid
5-HTP: 5-Hydroxytryptophan
5-HT: Serotonin
CaMKII: Calmodulin-dependent protein kinase II
EC: Enterochromaffin cells
GI: Gastrointestinal
IAA: 3-Indole acetic acid
IBS: Irritable bowel syndrome
IBS-D: Diarrhea-predominant irritable bowel syndrome
PA: Phenylacetic acid
PEA: Phenethylamine
Phe: Phenylalanine
PKA: Cyclic AMP-dependent protein kinase
TAAR1: Trace amine-associated receptor 1
TpA: Tryptamine
TPH1: Tryptophan hydroxylase 1
Trp: Tryptophan

## Acknowledgements

This work was kindly funded by the National Natural Science Foundation of China (82000504, 81973538), General Research Fund (12102620 and 12102721) and Innovation and Technology Fund (ITS/148/14FP). We are thankful to all patients and healthy volunteers who donated specimens for this study.

## Author contributions

ZX.B. and XHL.W conceptualized the project and designed the experiments. ZW.N. and LX.Z. conducted metabolites quantification in serum, urine and fecal samples of IBS-D patients. LX.Z., CH.H., YJ.Z. and M.Z. performed *in vivo* and *in vitro* study. W.Y contributed to the management and recourses of clinical specimens. XL.W contributed to bacterial culture and engineering. JJ.W. and EL.Z. contributed to metagenomic data analysis. LX.Z. and XHL.W analyzed data and wrote the manuscript with the input of co-authors. HT.X., L.Z., YY.L., CFW.C., JD.H., SF.Y. and KM.C. contribute to the study design, technical support and data analysis support towards this work.

## STAR Methods

### Reagent and resources sharing contact

Further information and requests for resources and reagents can be directed to and will be fulfilled by the Lead Contact Zhao-xiang Bian.

### Experimental models and subject details

#### Human study

To determine the changes of phenethylamine and 5-HT signaling in IBS-D patients, the human study was conducted as previously described (Zhao et al., 2019). Briefly, IBS-D patients (n=345) and healthy volunteers (n=91) were recruited following the required criteria in this previous study. Written consent was obtained from each subject before the collection of their specimens. Biological samples including serum, urine and feces of all subjects were collected and transported to the laboratory in dry ice and stored in −80 C. Details of clinical indexes and diagnostic data of all subjects can be found in Supplementary.

#### Mouse study

The mice study was approved by the Committee on the Use of Human & Animal Subjects in Teaching & Research at Hong Kong Baptist University (Hong Kong SAR, China). All experiments were performed under the regulation of the Animals (Control of Experiments) Ordinance of the Department of Health, Hong Kong SAR, China. Male C57BL/6 mice aged 6-8 weeks and weighed 20-25g were purchased from Laboratory Animal Services Centre, The Chinese University of Hong Kong (Hong Kong SAR, China) and raised in Animal Unit, School of Chinese Medicine, Hong Kong Baptist University. The mice were housed at a condition of 12 h light/dark cycle in a controlled temperature of around 25C with free access to food and water. The *in vivo* experiments were reported following ARRIVE guidelines (du Sert et al., 2020).

#### Organoids

Intestinal organoids were obtained from small intestines of 8-10 week old C57BL6 mice as previously described (Wong et al., 2019). Briefly, the mice were euthanized by carbon dioxide and their small intestine were isolated and flushed with ice-cold phosphate-buffered saline (PBS). The intestinal segments were obtained by longitudinal incision and incubated in Gentle Cell Dissociation Reagent (STEMCELL Technology) with gentle shaking at room temperature for 15 min. The intestinal segments were then filtered using a 70-μm cell strainer to get the cells of the intestinal crypts. About 500 crypts were grown in the matrix gel in Supplement with advanced DMEM/F12 medium and growth-factor-reduced Matrigel in a ratio of 1:3. The standard medium was replaced with advanced DMEM/F12 medium in Supplement with 2 mM Glutamax, 10 mM HEPES, 1mM N-acetyl-cysteine, B27 Supplement, N2 Supplement, recombinant murine EGF (50 ng/ml), recombinant human R-spondin 1 (500 ng/ml), and recombinant murine Noggin (50 ng/ml). For experiments evaluating the effects of phenethylamine on 5-HT level and EC cell differentiation, organoids were treated with/without phenethylamine at indicated dosages.

#### QGP-1 cells

QGP-1 cells were cultured in RPMI 1640 medium Supplement with 10% FBS. QGP-1 cells are a human pancreatic endocrine cell line that can produce 5-HT (Doihara et al., 2009). For experiments evaluating the effects of phenylalanine, phenethylamine and phenylacetic acid on 5-HT production, QGP-1 cells were treated with/without phenylalanine, phenethylamine and phenylacetic acid at indicated dosages and time.

#### Bacteria strains

*Ruminococcus gnavus* (strain ATCC 29149) was firstly grown on Tryptic soy broth (TSB) agar plate and cultured in TSB broth using single colony. *R. gnavus* (strain ATCC 29149) was collected from the medium by centrifuge at 3, 500 rpm for 10 min at room temperature. These bacteria strains were then prepared in 200 μL sterilized PBS and then delivered to pseudo germ-free mice by oral gavage. Fecal samples were collected daily for measurement of fecal phenethylamine levels in pseudo germ-free mice before and after oral administration of bacteria strains.

Phenethylamine-producing *E.coli* K12 was constructed using the tryptophan decarboxylase (TDC) gene (A7B1V0) from *R. gnavus*. The TDC gene was cloned into the vector and the resulting plasmid was transferred into *E.coli* K12 as previously described (Kelpšas and Wachenfeldt, 2019). The insertion of the TDC gene into the *E.coli* K12 was confirmed by PCR and *in vitro* phenethylamine production in LB broth containing 0.25% phenylalanine. A vector-only control strain of *E.coli* K12 was also constructed. These bacteria strains were also collected as previously described and prepared in 200 μL sterilized PBS and then delivered to pseudo germ-free mice by oral gavage.

### Study methods details

#### Pseudo germ-free mouse model

An broad-spectrum antibiotics mixture (ABX) containing vancomycin (100 mg/kg), neomycin (200 mg/kg), metronidazole (200 mg/kg) and ampicillin (200 mg/kg) was used to establish a pseudo-germ-free model in mice. Briefly, the antibiotics mixture was administered to mice by oral gavage for 10 consecutive days in prior to fecal microbial transplantation (FMT) or monocolonization study (Kennedy et al., 2018).

#### Metabolites quantification

An Agilent 1290 Infinity II UPLC system coupled to a triple quadrupole (QQQ) 6470 mass spectrometry was used for targeted metabolomics profiling study. A Waters BEH 2.1×50mm C18 1.7μm column with a pre-column was used. The mobile phase used in LC-MS-QQQ was A: water with 0.1% formic acid and B: acetonitrile with 0.1% formic acid. The gradients were set as 2% B (0-0.5 min), 2%-30% B (0.5-4 min), 30%-100% B (4-6 min), 100% B (6-8 min), 100%-2% B (8-8.1 min) and maintain in 2% B (8.1-10 min). The MS data were collected and processed by in-house software provided by Agilent. The standards list, MRM transition, fragmentor and collision energy and are listed in (Supplement Table.S3). Correlation analysis was conducted among serum 5-HT level, fecal PEA level and diarrhea-related symptoms in IBS-D patients. The spearman’s rank coefficient correlation analysis was used for correlations and the significant cut-off value was set at an FDR adjusted *p*-value <0.05.

#### Batch culture of fecal samples

About 50 mg fecal samples were mixed with 20x sterilized 1x PBS (m/v) and homogenized with tissuelyzer after adding steel beads. 20 μL fecal suspension from each sample was inoculated in 2mL TSB Supplemented with 0.25% Phe and incubated overnight under anaerobic conditions at 37C. After incubation, 100 μL medium was then used to determine Phe and PEA levels. Briefly, 400 μL MeOH was added to 100 μL medium and vigorously vortexed. After that, the mixture was centrifuged at 15, 000 rpm for 10 min at 4 C. 200 μL supernatant was used for LC-MS analysis.

#### Phenethylamine administration

To study phenethylamine effect on 5-HT production, phenethylamine at a dosage of 2mg/kg, 5mg/kg and 10mg/kg (dissolved in 0.5% CMC-Na) were administered to mice by oral gavage. After 15 min, mice treated with/without phenethylamine were sacrificed under isoflurane anesthesia and serum samples and fresh proximal colon and distal colon tissues were collected and stored at −80◻ until analysis.

#### *In vivo* measurements

Fecal pellet water content test: 1.5 mL Eppendorf tubes were pre-weighed and were used to collect fecal pellets from mice immediately after defecation. The tubes were then tightly closed to measure the wet weight. Afterward, the tubes were opened and placed in an oven at 60 C overnight to measure the dry weight. Fecal pellet water content was measured by subtracting dry weight from wet weight and normalizing it to the wet pellet mass. Fecal pellets from mice are also counted after defecation within 2 hours for defecation frequency test.

Carmine Red Assay: Mice were given 300 μL 6% carmine red solution (prepared by 0.5% methylcellulose) by oral gavage to measure the GI transit time after treatment with phenethylamine with indicated dosages. The mice were put on white sheet paper to track the red pellet in their cages after oral gavage. Total taken time for the appearance of the first red pellet after oral gavage was recorded as gut transit time.

#### Western blotting analysis

Colonic tissues were lysed in RIPA buffer with protease inhibitor cocktails in a tissuelyzer. The tissue lysates were then centrifuged at 15,000 rpm for 15 min at 4 ◻ and quantified by BCA kit for thier protein concentration. The normalized supernatant was mixed with 5x loading buffer and heated at 98 ◻ on a dry bath for 10 min. The samples were then determined according to the western blotting protocol provided by Abcam. The blots were incubated with HRP-linked anti-rabbit IgG or anti-mouse IgG and reacted with enhanced chemiluminescence.

#### Transplantation of human microbiota (FMT) in pseudo-germ-free mice

Fecal samples of HC and IBS-D subjects (n=8/group) were prepared as suspensions in PBS at a concentration of (100 mg/mL). ABX-treated pseudo germ-free mice were used as recipient of HC and IBS-D microbiota and were daily orally gavaged with human microbiota suspension at a dosage of 500mg/kg for 5 consecutive days. After FMT, GI transit, stool consistency and defecation frequency were measured. Fecal samples were collected at prior to FMT (after ABX intervention) and after FMT.

#### Phenethylamine-producing bacteria analysis in IBS-D patients

Metagenomics data of IBS-D patients and healthy volunteers were obtained as previously described (Han et al., 2021). Correlation analysis was conducted between serum 5-HT level and fecal PEA level with PEA-producing bacteria in IBS-D patients. The spearman’s rank coefficient correlation analysis was used for correlations and the significant cut-off value was set at an FDR adjusted p<0.05.

### Statistical analysis

Results were from multiple, at least three times independent experiments. Data were expressed as average and SD or SEM values of at least triplicate samples. Significance *p*-values were calculated using GraphPad Prism 8 and the *p*-value less than 0.05 was regarded as statistically significant. Wilcoxon rank-sum two-tailed test was used to determine metabolites difference between IBS-D and healthy subjects. Wilcoxon one-tailed test was used to determine IBS-D-associated gut microbiota produces higher phenethylamine in metagenomics data. Unpaired student’s t-tests or one-way ANOVA were used in other experiments as indicated.

## Supplemental information

**Supplement Figure.S1.**
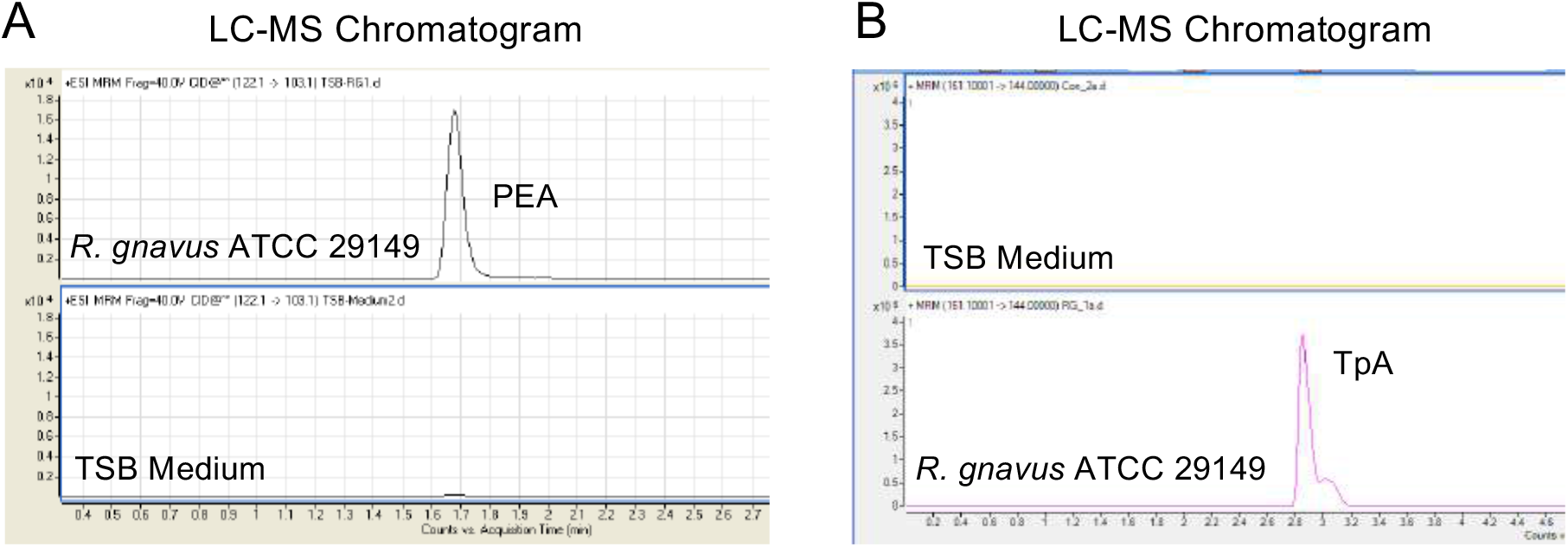
*R. gnavus* induces elevated 5-HT production and diarrhea-like symptoms. (A-B) LC-MS chromatogram of PEA and TpA level in tryptic soy broth culture medium of *R. gnavus* (ATCC 29149).

**Supplement Figure.S2.**
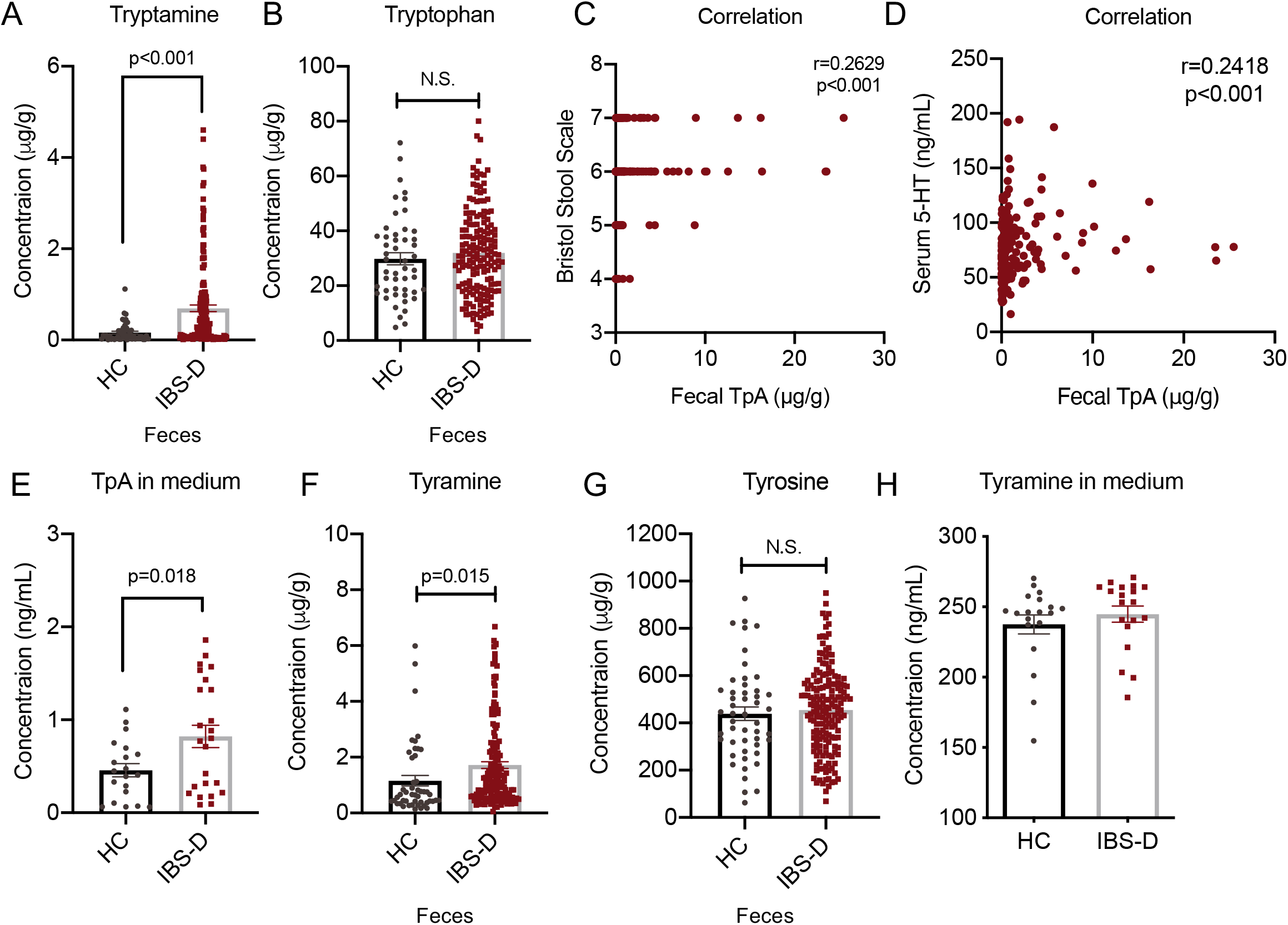
Fecal PEA and TpA are increased in IBS-D patients and positively correlated with serum 5-HT level. (A-B) Fecal TpA and Trp level in fecal samples of HC and IBS-D subjects. (C-D) Spearman’s correlation between fecal TpA with relative abundances of *R. gnavus* and Bristol Stool Scale in IBS-D subjects. (E) TpA level in batch culture samples using feces from HC and IBS-D (n=30/group). (F-G) Fecal tyramine and tyrosine level in HC and IBS subjects. (H) Tyramine level in batch culture samples using feces from HC and IBS-D (n=30/group). Differences of fecal TpA, Trp, tyramine and tyrosine level in HC and IBS subjects as well as batch culture samples were analyzed by one-tailed Mann-Whitney test. Data are presented as mean + S.E.M.

**Supplement Figure.S3.**
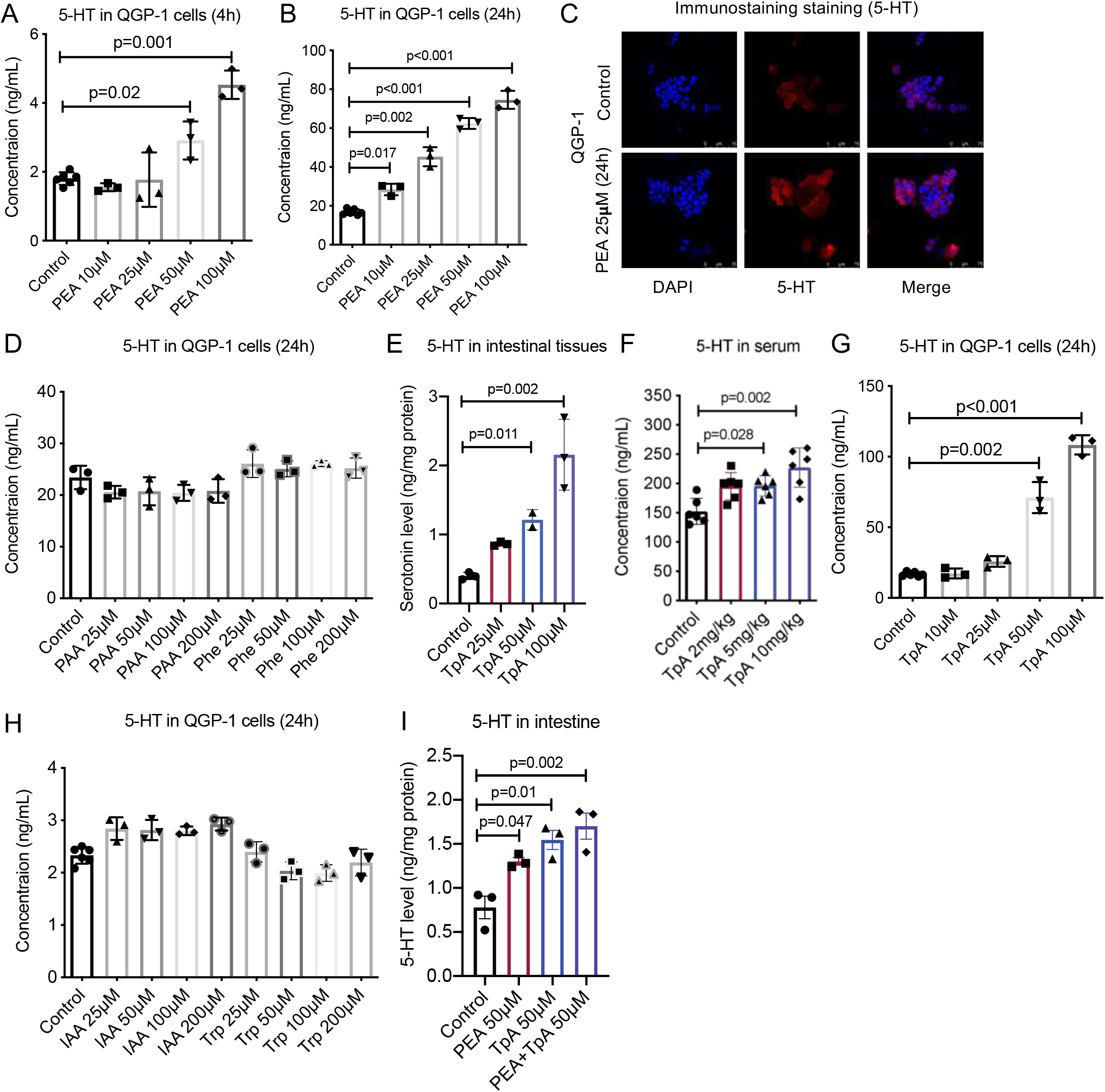
PEA and TpA activates 5-HT production, accelerates GI transit and increase colonic secretion *in vitro* and *in vivo*. (A-B) 5-HT level in QGP-1 cells after treatment of PEA as indicated concentration (25μM, 50μM and 100μM) or control for 4 hours and 24 hours (n=3/group). (C) IF staining of 5-HT in QGP-1 cells after treatment of PEA 25μM or control for 24 hours (n=3/group). (D) 5-HT level in QGP-1 cells after treatment of PAA or Phe as indicated concentration (50μM, 100μM and 200μM) or control for 24 hours (n=3/group). (E-F) 5-HT level in mice intestinal tissues and serum after treatment of TpA as indicated dosages (2mg/kg, 5mg/kg and 10mg/kg) or control (water) (n=6/group). (G) 5-HT level in QGP-1 cells after treatment of TpA as indicated concentration (25μM, 50μM and 100μM) or control for 24 hours (n=3/group). (H) 5-HT level in QGP-1 cells after treatment of IAA or Trp as indicated concentration (50μM, 100μM and 200μM) or control for 24 hours (n=3/group). (I) 5-HT level in mice intestinal tissues *ex vivo* after treatment of TpA and PEA as indicated dosages (50μM) (n=3/group). Differences of 5-HT in mice serum and intestine tissues and QGP-1 cells were analyzed by ordinary one-way ANOVA. Data are presented as mean + S.D.

**Supplement Figure.S4.**
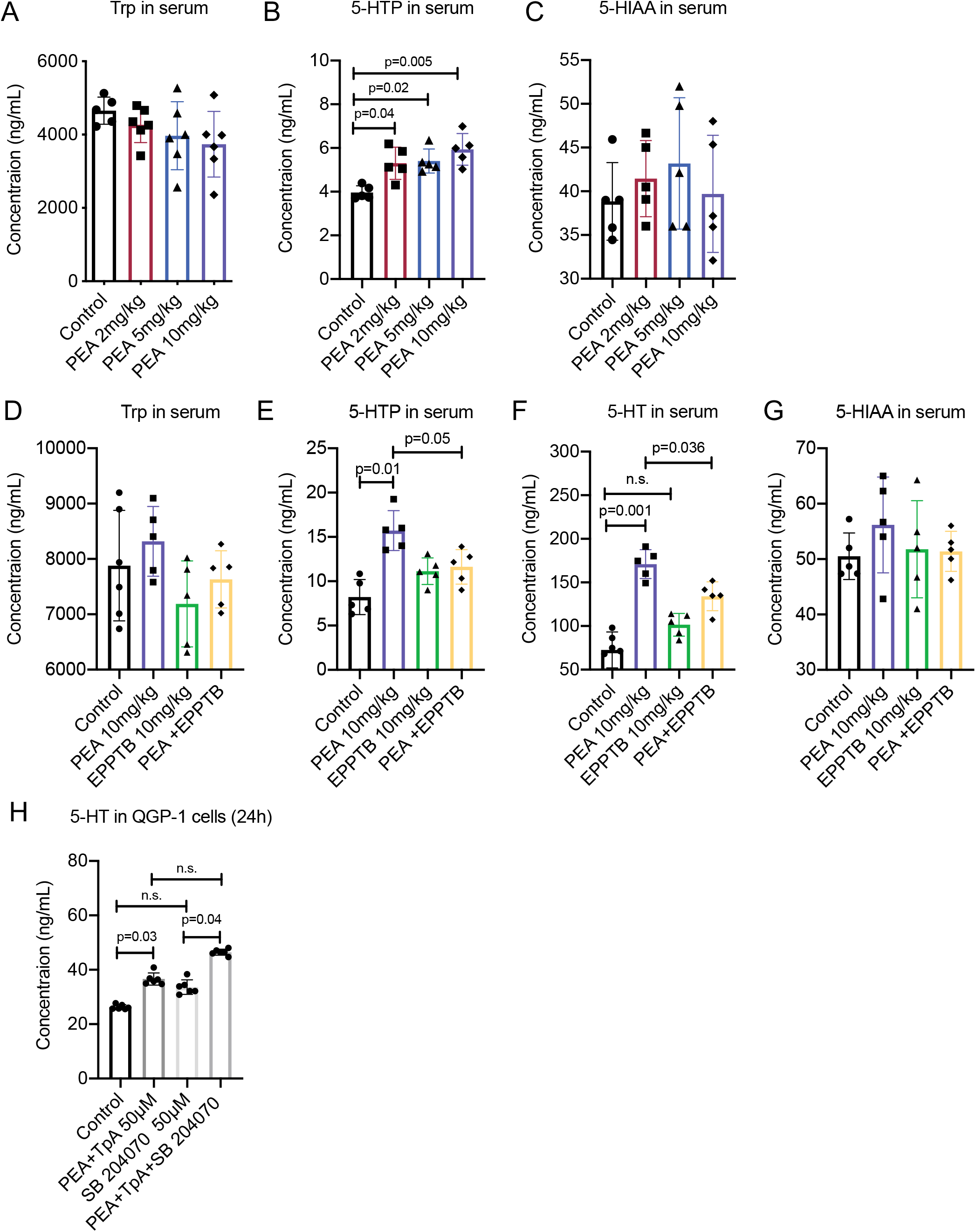
PEA stimulates 5-HT production via a TAAR1 dependent mechanism. (A-C) Serum Trp, 5-HTP and 5-HIAA in mice intestinal tissues and serum after treatment of TpA as indicated dosages (2mg/kg, 5mg/kg and 10mg/kg) or control (water) (n=6/group). (D-G) Serum Trp, 5-HTP, 5-HT and 5-HIAA in mice after treatment of PEA (10mg/kg) and TAAR1 antagonist EPPTB (10mg/kg) or control (1% DMSO in saline) (n=6/group). Differences of Trp, 5-HTP, 5-HT and 5-HIAA in mice serum and intestine tissues were analyzed by ordinary one-way ANOVA. Data are presented as mean + S.D.

**Supplement Figure.S5.**
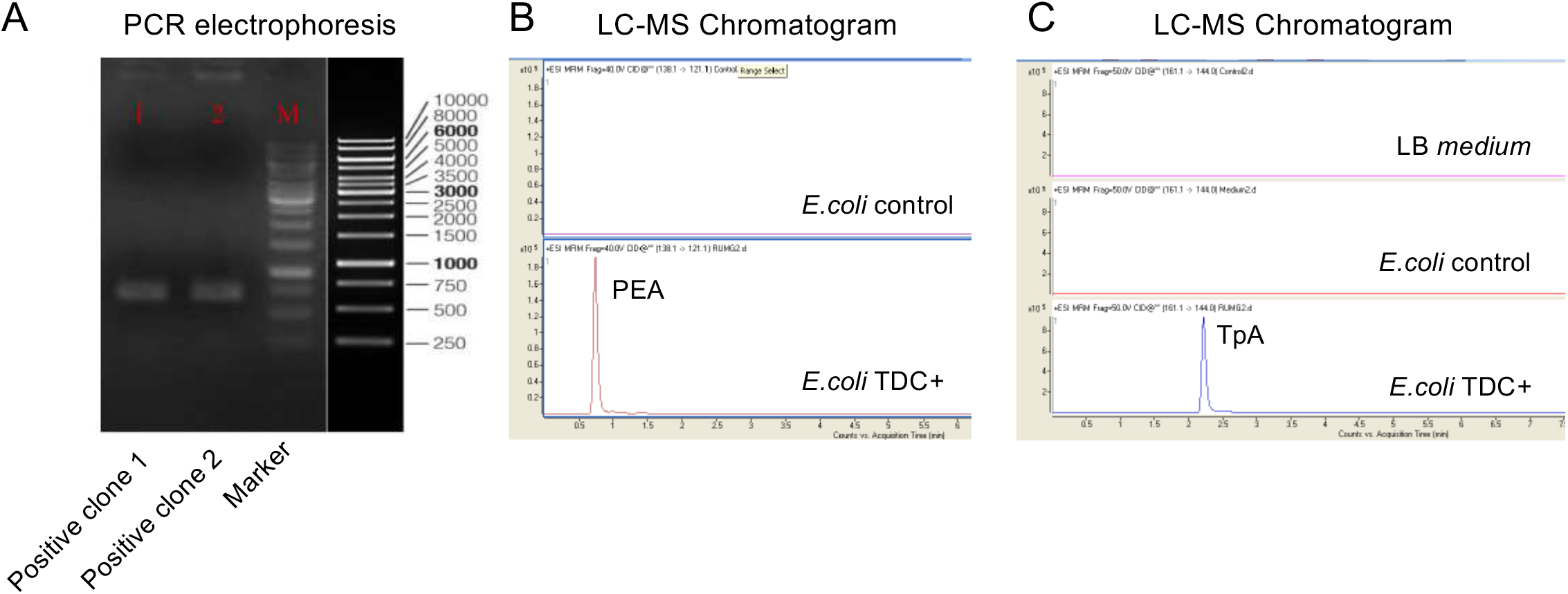
*In vivo* PEA and TpA produced by IBS-D associated bacteria enhances 5-HT synthesis and induce diarrhea-like symptoms. (A) PCR electrophoresis of positive clone of *E.coli* with TDC gene. (B-C) LC-MS chromatogram of PEA and TpA level in LB culture medium of *E.coli* TDC+ and *E.coli* vector control.

Supplement Table.S1 Spearman’s correlation between serum 5-HT with gut bacteria species in IBS-D patients.

Supplement Table.S2 Spearman’s correlation between fecal PEA with gut bacteria species in IBS-D patients.

Supplement Table.S3 MRM transition and parameters used for metabolites quantification.

Supplement Table.S4 Fecal PEA level in selected HC and IBS-D subjects for FMT experiment

## References

Agus, A., Planchais, J., and Sokol, H. (2018). Gut Microbiota Regulation of Tryptophan Metabolism in Health and Disease. Cell Host Microbe 23, 716–724.

Alnefeesi, Y., Tamura, J.K., Lui, L.M.W., Jawad, M.Y., Ceban, F., Ling, S., Nasri, F., Rosenblat, J.D., and McIntyre, R.S. (2021). Trace amine-associated receptor 1 (TAAR1): Potential application in mood disorders: A systematic review. Neurosci. Biobehav. Rev. 131, 192–210.

Bellono, N.W., Bayrer, J.R., Leitch, D.B., Castro, J., Zhang, C., O’Donnell, T.A., Brierley, S.M., Ingraham, H.A., and Julius, D. (2017). Enterochromaffin cells are gut chemosensors that couple to sensory neural pathways. Cell 170, 185–198.

Bhattarai, Y., Williams, B.B., Battaglioli, E.J., Whitaker, W.R., Till, L., Grover, M., Linden, D.R., Akiba, Y., Kandimalla, K.K., and Zachos, N.C. (2018). Gut microbiota-produced tryptamine activates an epithelial G-protein-coupled receptor to increase colonic secretion. Cell Host Microbe 23, 775–785.

Dodd, S., F. Carvalho, A., Puri, B.K., Maes, M., Bortolasci, C.C., Morris, G., and Berk, M. (2021). Trace Amine-Associated Receptor 1 (TAAR1): A new drug target for psychiatry? Neurosci. Biobehav. Rev. 120, 537–541.

Doihara, H., Nozawa, K., Kojima, R., Kawabata-Shoda, E., Yokoyama, T., and Ito, H. (2009). QGP-1 cells release 5-HT via TRPA1 activation; a model of human enterochromaffin cells. Mol. Cell. Biochem. 331, 239–245.

Duchemin, A., Berry, M.D., Neff, N.H., and Hadjiconstantinou, M. (2000). Phosphorylation and activation of brain aromatic L[amino acid decarboxylase by cyclic AMP[dependent protein kinase. J. Neurochem. 75, 725–731.

Dunlop, S.P., Jenkins, D., Neal, K.R., and Spiller, R.C. (2003). Relative importance of enterochromaffin cell hyperplasia, anxiety, and depression in postinfectious IBS. Gastroenterology 125, 1651–1659.

Fadgyas-Stanculete, M., Buga, A.-M., Popa-Wagner, A., and Dumitrascu, D.L. (2014). The relationship between irritable bowel syndrome and psychiatric disorders: from molecular changes to clinical manifestations. J. Mol. Psychiatry 2, 4.

Ge, X., Pan, J., Liu, Y., Wang, H., Zhou, W., and Wang, X. (2018). Intestinal crosstalk between microbiota and serotonin and its impact on gut motility. Curr. Pharm. Biotechnol. 19, 190–195.

Hall, A.B., Yassour, M., Sauk, J., Garner, A., Jiang, X., Arthur, T., Lagoudas, G.K., Vatanen, T., Fornelos, N., and Wilson, R. (2017). A novel Ruminococcus gnavus clade enriched in inflammatory bowel disease patients. Genome Med. 9, 1–12.

Han, L., Zhao, L., Zhou, Y., Yang, C., Xiong, T., Lu, L., Deng, Y., Luo, W., Chen, Y., Qiu, Q., et al. (2021). Altered metabolome and microbiome features provide clues in understanding irritable bowel syndrome and depression comorbidity. ISME J.

Henke, M.T., Kenny, D.J., Cassilly, C.D., Vlamakis, H., Xavier, R.J., and Clardy, J. (2019). *Ruminococcus gnavus*, a member of the human gut microbiome associated with Crohn’s disease, produces an inflammatory polysaccharide. Proc. Natl. Acad. Sci. 116, 12672 LP – 12677.

Van den Houte, K., Colomier, E., Schol, J., Carbone, F., and Tack, J. (2020). Recent advances in diagnosis and management of irritable bowel syndrome. Curr. Opin. Psychiatry 33, 460–466.

Jeffery, I.B., Das, A., O’Herlihy, E., Coughlan, S., Cisek, K., Moore, M., Bradley, F., Carty, T., Pradhan, M., Dwibedi, C., et al. (2020). Differences in Fecal Microbiomes and Metabolomes of People With vs Without Irritable Bowel Syndrome and Bile Acid Malabsorption. Gastroenterology 158, 1016–1028.e8.

Kelpšas, V., and Wachenfeldt, C. von (2019). Strain improvement of Escherichia coli K-12 for recombinant production of deuterated proteins. Sci. Rep. 9, 17694.

Kendig, D.M., and Grider, J.R. (2015). Serotonin and colonic motility. Neurogastroenterol. Motil. 27, 899–905.

Kennedy, E.A., King, K.Y., and Baldridge, M.T. (2018). Mouse microbiota models: comparing germ-free mice and antibiotics treatment as tools for modifying gut bacteria. Front. Physiol. 1534.

Kuhn, D.M., Arthur Jr, R., and States, J.C. (1997). Phosphorylation and activation of brain tryptophan hydroxylase: identification of serine◻58 as a substrate site for protein kinase A. J. Neurochem. 68, 2220–2223.

Kumer, S.C., Mockus, S.M., Rucker, P.J., and Vrana, K.E. (1997). Amino◻terminal analysis of tryptophan hydroxylase: protein kinase phosphorylation occurs at serine◻58. J. Neurochem. 69, 1738–1745.

Liu, Y., Hou, Y., Wang, G., Zheng, X., and Hao, H. (2020). Gut Microbial Metabolites of Aromatic Amino Acids as Signals in Host–Microbe Interplay. Trends Endocrinol. Metab.

Luo, M., Zhuang, X., Tian, Z., and Xiong, L. (2021). Alterations in short-chain fatty acids and serotonin in irritable bowel syndrome: a systematic review and meta-analysis. BMC Gastroenterol. 21, 14.

Marcobal, A., De Las Rivas, B., Landete, J.M., Tabera, L., and Muñoz, R. (2012). Tyramine and phenylethylamine biosynthesis by food bacteria. Crit. Rev. Food Sci. Nutr. 52, 448–467.

Mars, R.A.T., Yang, Y., Ward, T., Houtti, M., Priya, S., Lekatz, H.R., Tang, X., Sun, Z., Kalari, K.R., and Korem, T. (2020). Longitudinal multi-omics reveals subset-specific mechanisms underlying irritable bowel syndrome. Cell 182, 1460–1473.

Matthes, S., and Bader, M. (2018). Peripheral serotonin synthesis as a new drug target. Trends Pharmacol. Sci. 39, 560–572.

Neff, N.H., Duchemin, A., and Hadjiconstantinou, M. (2002). Activation of aromatic l[amino acid decarboxylase by calcium/calmodulin kinase II. J. Neurochem. 81, 98–99.

Öhman, L., and Simrén, M. (2010). Pathogenesis of IBS: role of inflammation, immunity and neuroimmune interactions. Nat. Rev. Gastroenterol. Hepatol. 7, 163–173.

De Palma, G., Lynch, M.D.J., Lu, J., Dang, V.T., Deng, Y., Jury, J., Umeh, G., Miranda, P.M., Pastor, M.P., and Sidani, S. (2017). Transplantation of fecal microbiota from patients with irritable bowel syndrome alters gut function and behavior in recipient mice. Sci. Transl. Med. 9.

Porter, C.K., Cash, B.D., Pimentel, M., Akinseye, A., and Riddle, M.S. (2012). Risk of inflammatory bowel disease following a diagnosis of irritable bowel syndrome. BMC Gastroenterol. 12, 55.

Rana, S. V, Sharma, S., Sinha, S.K., Parsad, K.K., Malik, A., and Singh, K. (2013). Pro-inflammatory and anti-inflammatory cytokine response in diarrhoea-predominant irritable bowel syndrome patients. Trop. Gastroenterol. 33, 251–256.

Reigstad, C.S., Salmonson, C.E., III, J.F.R., Szurszewski, J.H., Linden, D.R., Sonnenburg, J.L., Farrugia, G., and Kashyap, P.C. (2015). Gut microbes promote colonic serotonin production through an effect of short◻chain fatty acids on enterochromaffin cells. FASEB J. 29, 1395–1403.

Salahshouri, P., Emadi-Baygi, M., Jalili, M., Khan, F.M., Wolkenhauer, O., and Salehzadeh-Yazdi, A. (2021). A metabolic model of intestinal secretions: the link between human microbiota and colorectal cancer progression. Metabolites 11, 456.

Santoru, M.L., Piras, C., Murgia, A., Palmas, V., Camboni, T., Liggi, S., Ibba, I., Lai, M.A., Orrù, S., Blois, S., et al. (2017). Cross sectional evaluation of the gut-microbiome metabolome axis in an Italian cohort of IBD patients. Sci. Rep. 7, 9523.

du Sert, N.P., Hurst, V., Ahluwalia, A., Alam, S., Avey, M.T., Baker, M., Browne, W.J., Clark, A., Cuthill, I.C., and Dirnagl, U. (2020). The ARRIVE guidelines 2.0: updated guidelines for reporting animal research. BMJ Open Sci. 4, e100115.

Sotnikova, T.D., Caron, M.G., and Gainetdinov, R.R. (2009). Trace amine-associated receptors as emerging therapeutic targets. Mol. Pharmacol. 76, 229–235.

Sperber, A.D., Dumitrascu, D., Fukudo, S., Gerson, C., Ghoshal, U.C., Gwee, K.A., Hungin, A.P.S., Kang, J.-Y., Minhu, C., and Schmulson, M. (2017). The global prevalence of IBS in adults remains elusive due to the heterogeneity of studies: a Rome Foundation working team literature review. Gut 66, 1075–1082.

Stasi, C., Bellini, M., Bassotti, G., Blandizzi, C., and Milani, S. (2014). Serotonin receptors and their role in the pathophysiology and therapy of irritable bowel syndrome. Tech. Coloproctol. 18, 613–621.

Thijssen, A.Y., Mujagic, Z., Jonkers, D., Ludidi, S., Keszthelyi, D., Hesselink, M.A., Clemens, C.H.M., Conchillo, J.M., Kruimel, J.W., and Masclee, A.A.M. (2016). Alterations in serotonin metabolism in the irritable bowel syndrome. Aliment. Pharmacol. Ther. 43, 272–282.

Wei, W., Wang, H.-F., Zhang, Y., Zhang, Y.-L., Niu, B.-Y., and Yao, S.-K. (2020). Altered metabolism of bile acids correlates with clinical parameters and the gut microbiota in patients with diarrhea-predominant irritable bowel syndrome. World J. Gastroenterol. 26, 7153–7172.

Wong, H.L.X., Qin, H., Tsang, S.W., Zuo, X., Che, S., Chow, C.F.W., Li, X., Xiao, H., Zhao, L., and Huang, T. (2019). Early life stress disrupts intestinal homeostasis via NGF-TrkA signaling. Nat. Commun. 10, 1–14.

Xie, Z., and Miller, G.M. (2008). β-phenylethylamine alters monoamine transporter function via trace amine-associated receptor 1: implication for modulatory roles of trace amines in brain. J. Pharmacol. Exp. Ther. 325, 617–628.

Yano, J.M., Yu, K., Donaldson, G.P., Shastri, G.G., Ann, P., Ma, L., Nagler, C.R., Ismagilov, R.F., Mazmanian, S.K., and Hsiao, E.Y. (2015). Indigenous bacteria from the gut microbiota regulate host serotonin biosynthesis. Cell 161, 264–276.

Young, E.A., Neff, N.H., and Hadjiconstantinou, M. (1993). Evidence for cyclic AMP◻mediated increase of aromatic L◻amino acid decarboxylase activity in the striatum and midbrain. J. Neurochem. 60, 2331–2333.

Zhao, L., Yang, W., Chen, Y., Huang, F., Lu, L., Lin, C., Huang, T., Ning, Z., Zhai, L., and Zhong, L.L.D. (2019). A Clostridia-rich microbiota enhances bile acid excretion in diarrhea-predominant irritable bowel syndrome. J. Clin. Invest. 130.

Zhao, L., Yang, W., Chen, Y., Huang, F., Lu, L., Lin, C., Huang, T., Ning, Z., Zhai, L., and Zhong, L.L.D. (2020). A Clostridia-rich microbiota enhances bile acid excretion in diarrhea-predominant irritable bowel syndrome. J. Clin. Invest. 130, 438–450.

